# Emergence of preference coding in the macaque lateral prefrontal cortex by neurofeedback of unit activity related to working memory

**DOI:** 10.1101/2023.11.29.568968

**Authors:** Atsushi Noritake, Kazuyuki Samejima, Masataka Watanabe, Masamichi Sakagami

**Affiliations:** Division of Behavioral Development, Department of System Neuroscience, National Institute for Physiological Sciences, National Institutes of Natural Sciences; Department of Physiological Sciences, School of Life Science, The Graduate University for Advanced Studies (SOKENDAI); Tamagawa University, Brain Science Institute; Tokyo Metropolitan Institute of Medical Sciences

**Keywords:** Neurofeedback, working memory, lateral prefrontal cortex, macaque monkey, single-unit recording

## Abstract

Techniques utilizing neurofeedback, a form of biofeedback using neural signals from the brain, have been applied lately to higher association areas such as the lateral prefrontal cortex (LPFC); however, it remains unexplored how well neurofeedback using unit activity in the LPFC modulates its working memory-related activity and performance. To address this issue, we provided neurofeedback of LPFC unit activity during a delay period to two monkeys while they performed a delayed matching-to-paired-sample task. In the task, neurofeedback allowed the animals to shorten the delay length by increasing delay activity and make an earlier choice. Neurofeedback significantly increased delay activity in two-thirds of task-related neurons. Notably, in 16% of these neurons, a preference for delay activity and performance dependent on the stimulus emerged. Although neurofeedback decreased performance primarily due to choice errors, the disassociation of neurofeedback linkage rescued performance. Further, the neuronal activity of simultaneously recorded neurons without neurofeedback linkage suggests that neurofeedback reconfigured the net activity of the LPFC to adapt to new situations. These findings indicate that LPFC neurons can dynamically multiplex different types of information to adapt to environmental changes. Thus, we demonstrated the significant potential of neurofeedback using unit activity to investigate information processing in the brain.

## Introduction

Working memory (WM), which is highly developed in primates, maintains relevant information and suppresses irrelevant information over time. WM enables animals to adopt a consistent strategy and behavior to adapt to environmental changes. Previous studies revealed that neuronal activity in the lateral prefrontal cortex (LPFC) is profoundly involved in WM function (Parthasarathy et al., 2017; Rouzitalab et al., 2023; Sakagami and Niki, 1994; Spaak et al., 2017). Electrophysiological analysis in non-human primates has shown that after training in WM tasks, the number of recruited PFC neurons increases during stimulus presentation and a delay period (Qi et al., 2011, 2012). Moreover, the variability of firing rates across trials and correlation discharges, such as noise correlation and across-trial correlation between neurons, decreases (Qi and Constantinidis, 2012a, b). These findings shed light on PFC neural plasticity and information-coding manner of the PFC in which neurons can tune their activity to adapt to environmental changes through training (Duncan, 2001; Miller and Cohen, 2001). However, studying such learning via training requires devices to track and monitor long-term changes in neural activity, and it remains unclear how such modifications occur.

Feedback techniques using biological signals, termed biofeedback, such as myopotentials (Giggins et al., 2013), electroencephalograms (Gruzelier, 2014), and brain blood oxygen level-dependent signals (Shibata et al., 2011), have a significant potential by quickly making a shortcut link between activity, behavior, and outcome, e.g., reward or punishment. In particular, neurofeedback, a biofeedback technique using neural signals from the brain, has been utilized preferentially to target neural activity in motor-related brain areas (Andersen et al., 2004; Carp et al., 2006; Chapin et al., 1999; Fetz and Baker 1973) including the primary motor area, premotor areas, and frontal eye field. Yet, few studies have examined how neurofeedback modulates activity in higher association areas related to cognitive functions (Cerf et al., 2010; Ishikawa et al., 2014), such as the LPFC (Seitz, 2013; Zhang et al., 2013). A pioneering study demonstrated that visual neurofeedback of single-unit activity in the primate LPFC could increase or decrease its activity in a task-dependent manner (Kobayashi et al., 2010). In that study, the animals could move visual stimuli on a display using LPFC unit activity and attain a liquid reward when the increased or decreased neuronal activity reached some criteria, i.e., neuronal modulation resulted in reward delivery. However, as many LPFC neurons are thought to encode WM-related information (Amemori and Sawaguchi, 2006; Kennerley and Wallis, 2009; Kobayashi et al., 2002; Leon and Shadlen, 1999; Rainer et al., 1999), neurofeedback of their activity could be more directly associated with aspects of WM, rather than reward and its expectation (Lauwereyns et al., 2001; Watanabe et al., 2007). For example, neurofeedback using delay activity, which is not directly associated with outcome, may modify the WM component and improve or impede WM task performance. In addition, neurofeedback allows us to track unit activity modulation more easily through environmental changes, i.e., before and after neurofeedback, compared with long-term training.

To explore this possibility, we provided neurofeedback of single-unit activity in the LPFC only during a delay period while monkeys performed a delayed matching-to-paired-sample task (DMPST) (Rainer et al., 1999; Sakai and Miyashita, 1991). In this task, the animal was required to choose the paired associate of a sample stimulus between options after a delay period for a liquid reward (**Fig. 1**). We provided visual feedback of neuronal activity (neurofeedback) in the LPFC during the delay period (**Fig. 1b,d**) and examined how neurofeedback modified delay activity for two monkeys. Neurofeedback allowed the animals to shorten the length of the delay period by increasing firing rates during the delay period and have an earlier chance of making a choice. Conversely, the delay period could be lengthened when neuronal delay activity was decreased. Thus, neurofeedback modification of delay activity did not seem to be linked directly to reward-related components, but rather to WM components.

**Figure 1.**
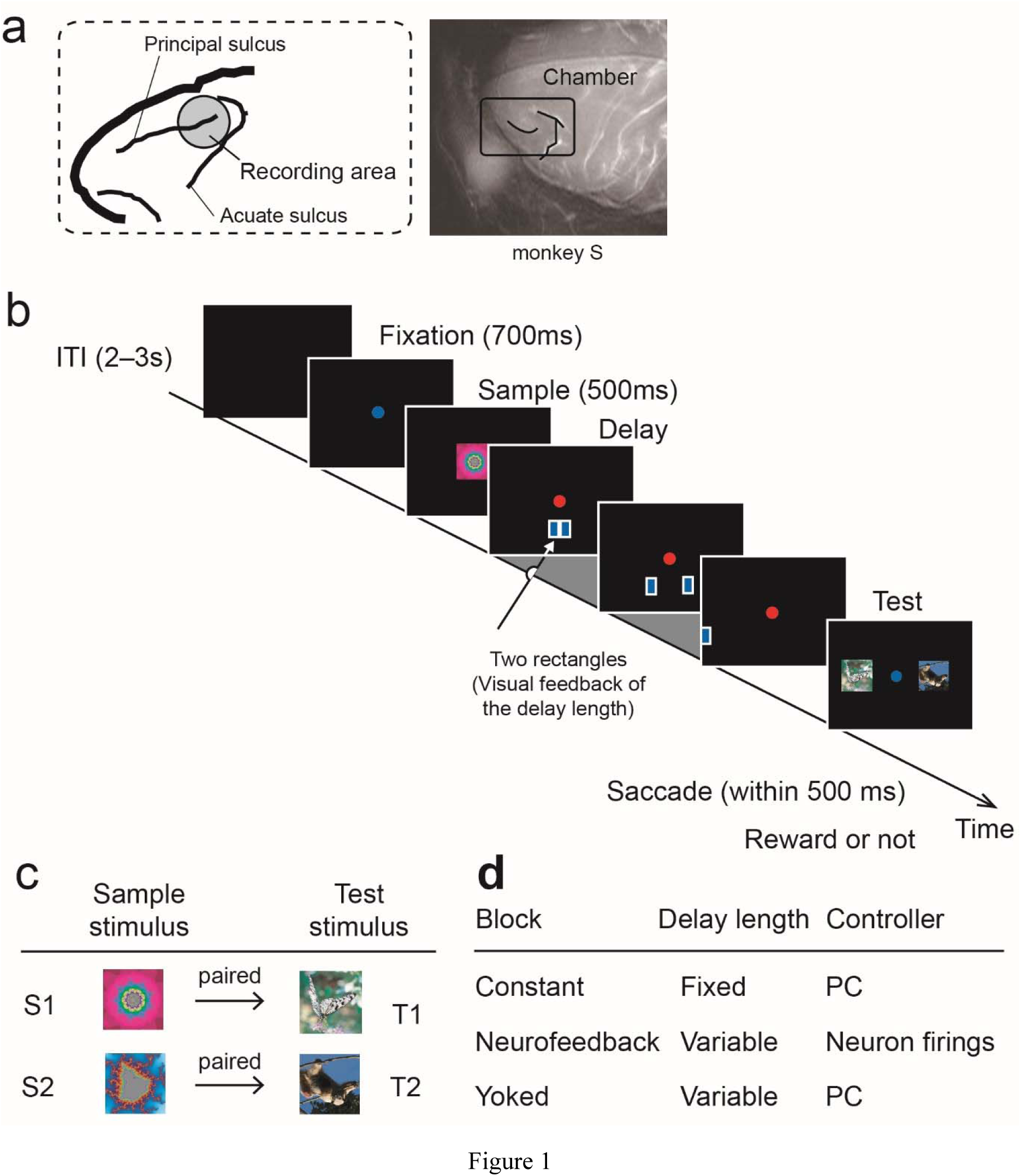
Recording area and delayed matching-to-paired-sample task (DMPST). **a**. Schematic cortical map of the recording site in the left lateral prefrontal cortex (*left*) and magnetic resonance image of monkey S (*right*). Neurons around the caudal part of the principal sulcus were recorded (shaded circle area). **b.** Task sequence of behavioral events. Gray area depicts the delay period. ITI, inter-trial interval **c.** Sample–test stimulus matching in the DMPST. **d.** Block sequence, delay-length modulation, and the controller of rectangle movement during the delay period.

We hypothesize that neurofeedback motivated the animals to shorten the length of the delay period and improve performance by increasing neuronal activity in the LPFC predominantly during the delay period because previous studies on WM function revealed that monkeys perform better as delay length decreases (e.g., Sakurai, 2001).

## Results

We recorded the unit activity of neurons in the left LPFC of two monkeys (**Fig. 1a**) while they performed a DMPST (**Fig. 1b,c**) with and without neurofeedback (see Methods; **Fig. 1d**) in two trial blocks: constant and neurofeedback. During the delay period in the task, two rectangles appeared at the center of a screen and moved horizontally, away from each other, toward the right and left edges of the screen at the same speed (**Fig. 1b,c**). They disappeared when they reached the edges. In the constant blocks, the movement of the rectangles was constant and not associated with delay activity. In the neurofeedback blocks, their movement was associated with delay activity, and therefore, the animals could shorten or lengthen the delay length by increasing or decreasing the firing rate during the delay period, respectively. Accordingly, rectangle movement speed and delay length varied across trials. Constant blocks always preceded neurofeedback blocks to determine the step size, i.e., the speed, of rectangles (see Methods). A pair of the constant and neurofeedback blocks was referred to as a session. In addition, another control block, a “yoked” block, was introduced to the animals after some of the sessions (**Fig. 1d**). In the yoked blocks, the association between delay activity and length was dissected under the control of delay-length variance among trials by replaying the same sequence of delay lengths experienced in the preceding neurofeedback blocks. In contrast to the variable step timing of rectangle movement during the neurofeedback blocks, the rectangles moved at a constant speed in the yoked blocks.

### Neuronal and behavioral modulation by neurofeedback

We conducted 140 sessions (monkey M: 39 sessions; monkey S: 101 sessions), i.e., 140 neurons were tested under neurofeedback. Among them, we further analyzed 125 neurons that exhibited significantly modulated activity during the constant blocks in the DMPST (task-related activity; see Methods; monkey M: 37 sessions; monkey S: 88 sessions). To assess how neuronal activity was modulated by neurofeedback and the stimuli, we applied two-way analysis of variance (ANOVA) with neurofeedback (with or without neurofeedback) and stimulus (S1 or S2) as factors to the firing rate during the delay period for each neuron. Note that in this study, the firing rate was expressed in spikes/s, meaning that in the neurofeedback blocks, the spike number during the delay period was divided by the delay length varying among trials. As a result, neurofeedback significantly increased and decreased the activity of 82 (65.6%, increased type) and 9 (7.2%, decreased type) neurons, respectively, during the delay period (neurofeedback: *p* < 0.05, two-way ANOVA) compared to the constant blocks. To test whether these neurons modulated their activity predominantly during the delay period, we applied repeated one-way ANOVA to the following activity: pre-sample activity (-500–-201 ms before sample onset), post-sample activity (201–500 ms after sample onset), and delay activity (201–500 ms after delay onset) with the Tukey-Kramer test as a *post-hoc* test. For the increased type, the delay activity was significantly more than the other two tested activities (*p* = 6.4 × 10^-3^, *F*_[2, 162]_ = 5.2, repeated one-way ANOVA; *p* < 0.05, Tukey-Kramer test). For the decreased type, the modulation of delay activity was the largest among the three epochs compared, but it was not significant (*n.s.*, repeated one-way ANOVA). The remaining 34 neurons (27.2%) were not modulated significantly (non-modulated type; neurofeedback, *n.s.*, two-way ANOVA). Thus, the animals shortened the delay length (**Fig. 2a**; median: 235 ms, *p* = 5.6 × 10^-7^, *Z* = 5.0, Mann-Whitney *U*-test) by increasing the delay activity of most of the tested neurons.

**Figure 2.**
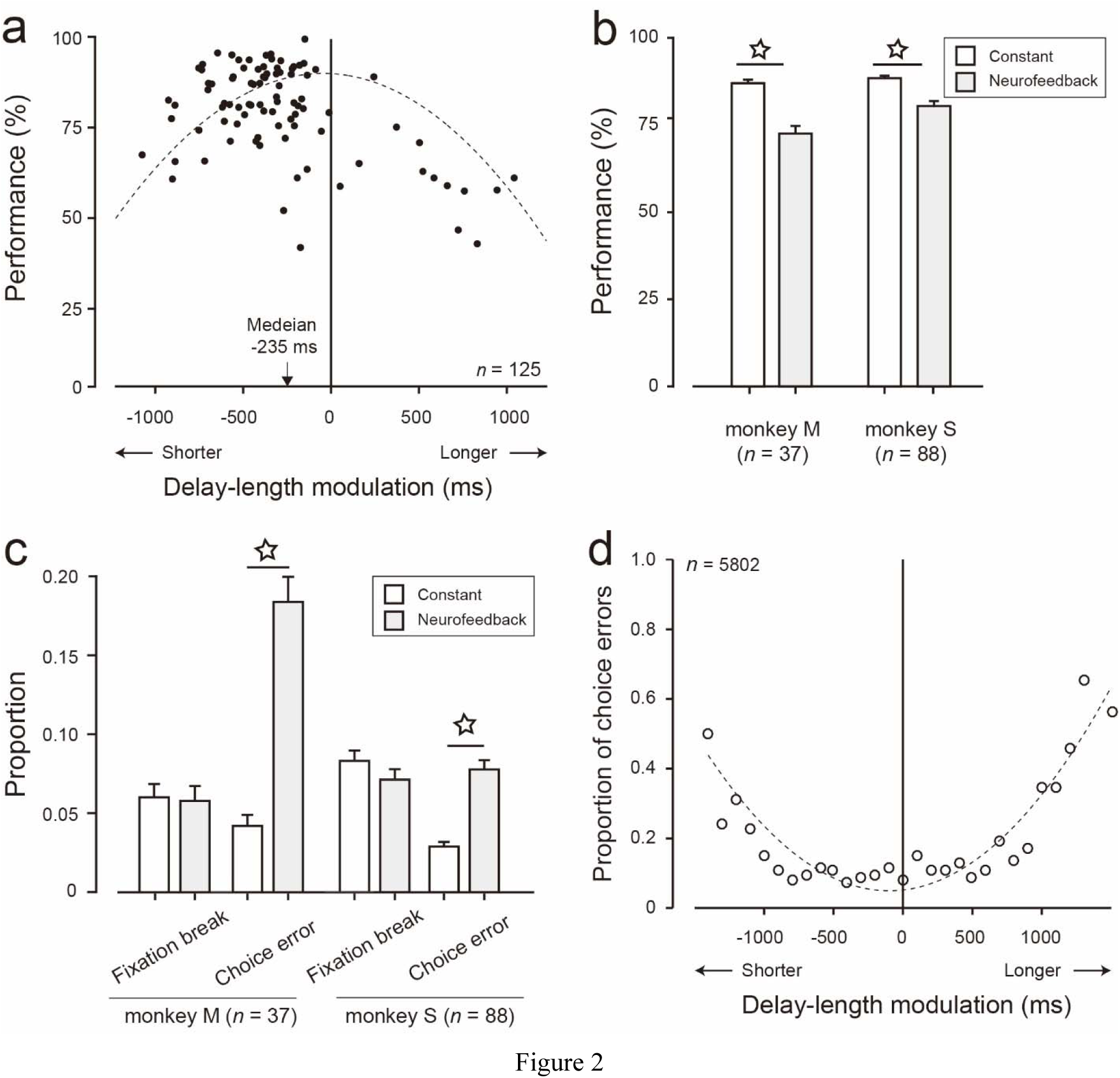
Performance of delayed matching-to-paired-sample task with and without neurofeedback. **a.** Shorter delay lengths in the neurofeedback blocks than in the constant blocks (*p* < 0.01, Mann-Whitney *U*-test) with decrease in correct performance during the excessive delay-length modulation. Downward arrow indicates the median delay-length modulation in the neurofeedback blocks. Solid vertical line represents zero delay-length modulation, corresponding to the delay length in the constant blocks. Dashed curve depicts the fitted quadratic regression curve. **b.** Correct performance (monkey M, *left*; monkey S, *right*) in the constant (white bars) and neurofeedback (gray bars) blocks. **c.** Proportion of error types (fixation-break and choice errors) in the constant (white bars) and neurofeedback (gray bars) blocks. **d.** Choice-error proportion as a function of delay-length modulation. The mean proportion of choice error was calculated in each bin (100 ms, no overlap) of the delay-length modulation. The same convention was used in **b** and **c**; white stars denote significant differences (*p* < 0.01, paired *t*-test). Error bars represent the standard error of the mean.

Further, we assessed another 54 simultaneously recorded neurons, which exhibited task-related activity, without neurofeedback linkage (“irrelevant” neurons) to determine whether our neurofeedback method using single-unit activity affected recorded activity specifically or background net activity of neurons simultaneously in the LPFC. In these neurons, 30 (55.6%) were increased type, and 11 (20.2%) were decreased type; the remaining 13 neurons (24.2%) were non-modulated type. These findings indicate that neurofeedback could increase the delay activity of more than 50% recorded irrelevant neurons, suggesting that neurofeedback modulated LPFC net activity.

Such neuronal modulation by neurofeedback decreased the correct performance rate compared to that in the constant blocks for both animals (monkey M: 87.4% in the constant blocks vs. 73.3% in the neurofeedback blocks, *p* = 3.3 × 10^-7^, *Z* = 5.1; monkey S: 88.4% in the constant blocks vs. 80.4% in the neurofeedback blocks, *p* = 5.7 × 10^-5^, *Z* = 4.0, Mann-Whitney *U*-test; **Fig. 2b**). The decrease in correct performance was dominated by an increase in the choice-error rate (monkey M: *p* = 4.4 × 10^-10^, *Z* = 6.2, monkey S: *p* = 3.2 × 10^-13^, *Z* = 7.3, Mann-Whitney *U*-test; **Fig. 2c, *right two bars for each monkey***) because the fixation-break error rate did not increase in either monkey (**Fig. 2c, *left two bars for each monkey***). Further, the decrease in correct performance (**Fig. 2a**) and the increase in choice error (**Fig. 2d**) occurred frequently in parallel when the modulation of delay length in the neurofeedback blocks relative to the delay length in the constant blocks (“delay-length modulation,” defined by subtracting the delay length in the neurofeedback blocks from that in the preceding constant blocks) was large. We plotted the choice-error (**Fig. 2d**) rates as a function of delay-length modulation to visualize this tendency. The correct performance rate was greater than 75% in most of the neurofeedback blocks when the modulation of delay length was moderate (**Fig. 2a**; moderate delay-length modulation: -600–0 ms), but decreased when the deviation of delay-length modulation became larger, indicating an inverted U-shape function in the correct performance rate against delay-length modulation during the neurofeedback blocks (*p* = 2.0 × 10^-10^, *F*_[2, 122]_ = 26.9, quadratic regression test). Concurrently, the choice-error rate was <10% when the modulation of delay length was moderate (-600–0 ms), while it gradually increased for the other delay-length modulations, indicating a U-shape function (**Fig. 2d**; *p* = 7.7 × 10^-10^, *F*_[2, 25]_ = 54.5, quadratic regression test). These results suggest that the larger deviation in neuronal modulation by neurofeedback that increased delay-length modulation resulted in lower correct performance and more choice errors.

### Saccadic latency in the neurofeedback blocks

When delay length became short, the animal’s decision during neurofeedback might become reactive and immature due to the sudden termination of the delay period, introducing reactive responses (i.e., short saccadic latency) and choice errors. However, saccadic latency at choice errors in such a situation (delay-length modulation: <-600 ms), as an index of reactive and immature decision making, was similar (mean: 215.3 ms) to that in the correct trials under neurofeedback (mean: 218.9 ms; *p* = 1.9 × 10^-1^, *t_211_* = 1.3, Welch’s *t*-test) as well as in the choice-error trials in the constant blocks (mean: 211.9 ms; *p* = 3.2 × 10^-1^, *t_356_*= 1.0, Welch’s *t*-test). In contrast, when delay length became long, saccadic latency was significantly longer than that in the choice-error trials in the constant blocks (mean saccadic latency: 222.4 ms in the constant blocks and 226.2 ms in the neurofeedback blocks; *p* = 1.2 × 10^-2^, *t_178_* = 2.5, Welch’s *t*-test) and in the correct trials in the neurofeedback blocks (mean: 211.9 ms; *p* = 4.2 × 10^-5^, *t_303_* = 2.5, Welch’s *t*-test). Thus, choice errors were induced without reactive and immature responses in extensively short delay periods and with more extended saccadic responses in extensively long delay periods.

### Correct performance in the yoked blocks

It is feasible that the neuronal modulation induced by neurofeedback modulated performance in the neurofeedback blocks. However, the variance of delay-length modulation might also affect correct performance in the neurofeedback blocks. In fact, the variance of delay-length modulation among trials was negatively correlated with correct performance in the neurofeedback blocks (stimulus S1: *r* = -0.25, *p* = 5.5 × 10^-3^; stimulus S2: *r* = -0.37, *p* = 2.5 × 10^-5^, Pearson’s correlation test; **Fig. 3a**). To assess the possibility that the variance of delay-length modulation caused the decrease in correct performance, we introduced another control block (yoked block) to the animals after the neurofeedback blocks in 95 of 125 sessions (15 sessions for monkey M and 80 sessions for monkey S). In the yoked blocks, the association between delay activity and length was dissected under the control of delay-length variance among trials.

**Figure 3.**
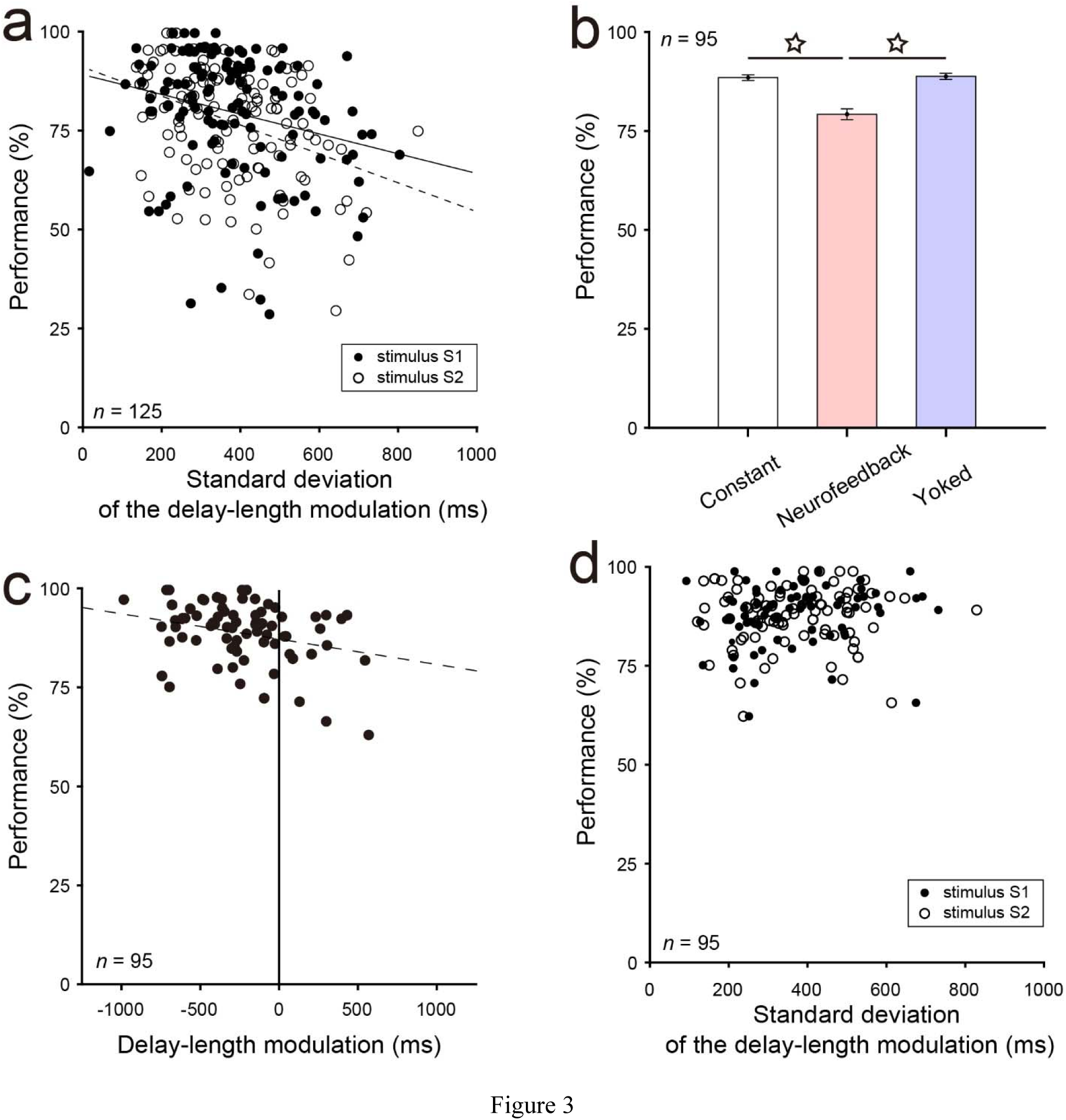
Performance in the yoked blocks compared with the constant and neurofeedback blocks. **a.** Larger standard deviation in the neurofeedback blocks induced lower correct performance. Solid and dashed lines indicate the significance of linear regression analysis (*p* < 0.01, Pearson’s correlation test) for stimuli S1 and S2, respectively. **b.** The decreased correct performance in the neurofeedback blocks recovered in the yoked blocks to the same level as in the constant blocks. Error bars represent the standard error of the mean. White stars indicate significant differences (*p* < 0.01, Tukey-Kramer test). **c.** Correct performance was not well fitted with a quadratic regression curve, but instead was well fitted with a linear regression line (dashed line). **d.** The same trial variance of the delay length used in the neurofeedback blocks did not correlate with the correct performance for either stimulus in the yoked blocks.

Correct performance in the yoked blocks recovered to a similar level as that in the constant blocks (88.5% in the constant blocks; 79.2% in the neurofeedback blocks; 88.8% in the yoked blocks; **Fig. 3b**), confirmed by one-way ANOVA (*p* = 2.3 × 10^-12^, *F*_[3, 282]_ = 29.5) with the Tukey-Kramer test as a *post-hoc* test (*p* = 0.97 between the constant and yoked blocks; *p* = 1.8 × 10^-10^ between the constant and neurofeedback blocks; *p* = 4.0 × 10^-11^ between the neurofeedback and yoked blocks). This recovery of performance was primarily due to the decrease of the choice error proportion relative to that in the neurofeedback blocks (yoked blocks vs. neurofeedback blocks: 4.4% vs. 8.6%, respectively, *p* = 7.9 × 10^-16^, *t*_139_ = 9.1, paired *t*-test). To evaluate performance in the yoked blocks further, we plotted the correct performance rate in each session as a function of delay-length modulation (**Fig. 3c**). Regression analysis revealed that performance was fitted in a linear, but not quadratic, manner (*p* = 6.6 × 10^-3^, *r* = -2.8 × 10^-2^, Pearson’s correlation test). Moreover, correct performance was not significantly influenced by the variance of delay-length modulation among the trials for each stimulus (stimulus S1: *p* = 7.7 × 10^-2^, *r* = 1.8 × 10^-1^; stimulus S2: *p* = 3.4 × 10^-1^, *r* = 9.9 × 10^-2^, Pearson’s correlation test; **Fig. 3d**). Thus, the delay-length variance among trials in the neurofeedback blocks was not the primary cause of the decrease in correct performance, suggesting that the neuronal modulation induced by neurofeedback during the delay period influenced performance.

### Three types of LPFC neural-code modulations by neurofeedback

On the basis of the results of two-way ANOVA, we further classified the increased and decreased types separately into three neuronal subtypes: Type I, II, and III. Type I neurons showed a significant effect of neurofeedback only (neurofeedback, *p* < 0.05; stimulus, *n.s.*; interaction between the neurofeedback and stimulus factors, *n.s.*; **Fig. 4a, *top***). Type II neurons exhibited significant neurofeedback and stimulus effects (neurofeedback, *p* < 0.05; stimulus, *p* < 0.05; interaction, *n.s.*; **Fig. 4a, *middle***). Type III neurons exhibited a significant neurofeedback effect with interaction (neurofeedback, *p* < 0.05; interaction, *p* < 0.05; **Fig. 4a, *bottom***), irrespective of the stimulus.

**Figure 4.**
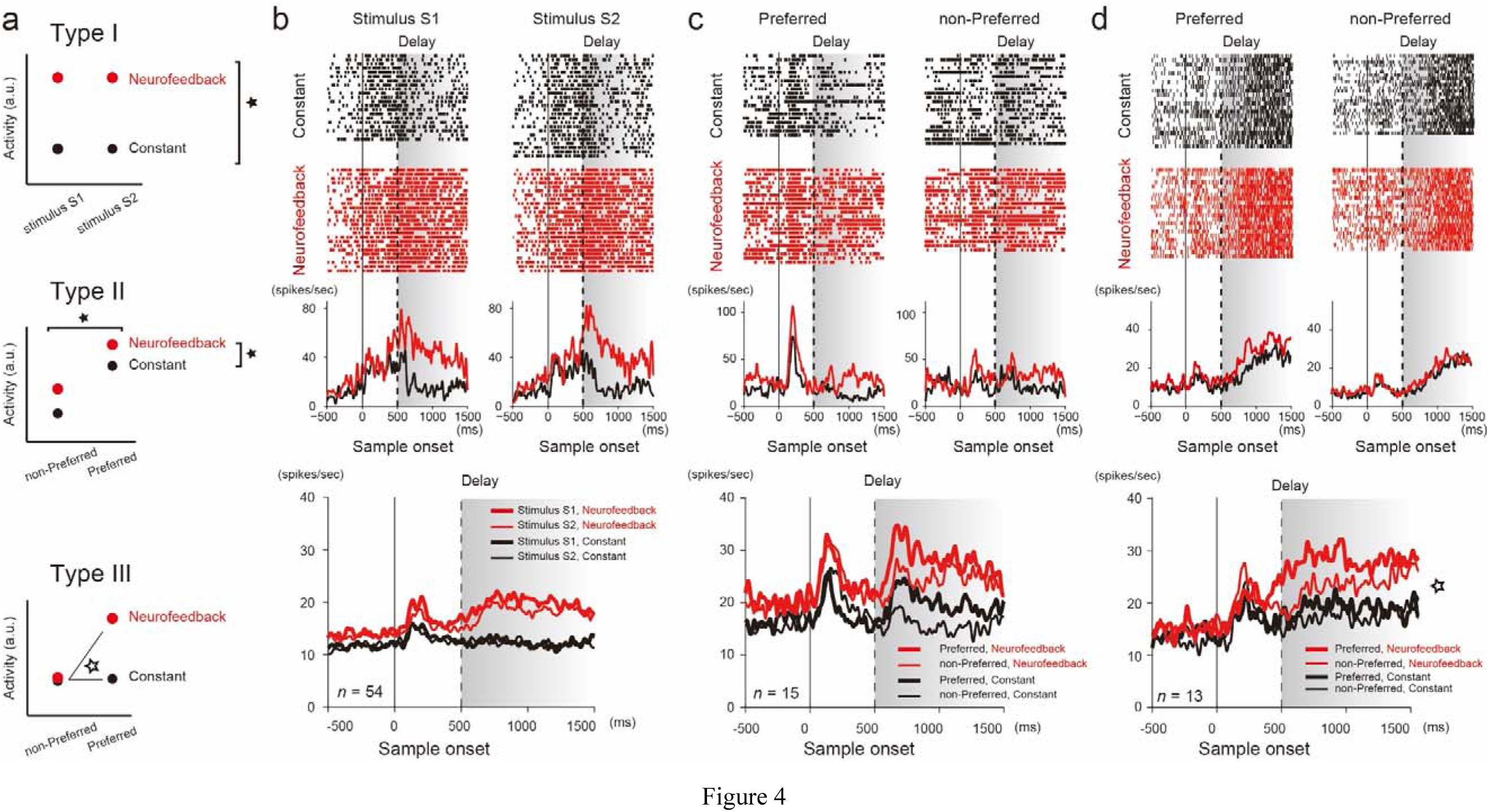
Activity of the modulated neuronal types. **a.** Schematic activity of Type I (*upper*), II (*middle*), and III (*bottom*) neurons. Filled and open stars indicate the statistical significance of factor effects and their interaction, respectively. a.u. arbitrary units. **b–d.** Activity of Type I (**b**), II (**c**), and III (**d**) neurons. Representative (*upper*) and population (*bottom*) activities of each neuronal type are displayed. Rastergrams and histograms of the representative neurons are aligned to the time of sample stimulus onset. Black and red tick marks depict spike timing in the constant and neurofeedback blocks, respectively. Vertical dashed lines indicate the start of the delay period. Curves illustrate the mean firing rates of population activity (Type I: *n* = 54; Type II: *n* = 15; Type III: *n* = 13) for the different sample stimuli (Type I: stimulus S1, thin curves; stimulus S2, thick curves; Type II and III: non-preferred stimulus, thin curves; preferred stimulus, thick curves). Shaded areas with gradation represent variable delay duration. Note that the preferred and non-preferred stimuli were determined dependent on the delay activity in the neurofeedback blocks. Colors represent the constant (black) and neurofeedback (red) blocks.

For the increased type neurons, approximately two-thirds of them (54/82, 65.9%) were Type I (example, **Fig. 4b, *top***; population, **Fig. 4b, *bottom***); these neurons increased their activity predominantly in the delay period, irrespective of the sample stimulus, during the neurofeedback blocks compared to the constant blocks. Fifteen neurons (18.3%) were Type II (example, **Fig. 4c, *top***; population, **Fig. 4c, *bottom***); these neurons had significantly increased delay activity and preserved the differences between the sample stimuli during neurofeedback. The sample stimulus that induced more delay activity in the neurofeedback blocks was referred to as the “preferred” stimulus and the other as the “non-preferred” stimulus. A relatively small number of neurons were Type III (13/82, 15.8%; example, **Fig. 4d, *top***; population, **Fig. 4d, *bottom***); these neurons exhibited similar delay activity between the sample stimuli during the constant blocks. However, in the neurofeedback blocks, a significant difference in delay activity emerged between the sample stimuli. The sample stimulus that induced more delay activity in the neurofeedback blocks was also referred to as the “preferred” stimulus and the other as the “non-preferred” stimulus. All neuronal types exhibited significantly increased activity for both stimuli during the sample period (1–500 ms after stimulus onset) in the neurofeedback blocks compared to the constant blocks (*p* < 0.01, paired *t*-test). In addition, only Type I neurons also showed significantly increased activity for both stimuli before stimulus onset (1–500 ms before stimulus onset in the fixation period; stimulus S1: *p* = 2.1 × 10^-4^, *t*_53_ = 4.0; stimulus S2: *p* = 3.0 × 10^-5^, *t*_53_ = 4.8). For the decreased type neurons, nine, five, and one were Type I, II, and III, respectively. These neuronal types did not exhibit significantly decreased activity for either stimulus during the fixation and sample periods in the neurofeedback blocks compared to the constant blocks (*n.s.*; **Fig. S1**), except for stimulus S1 during the pre-sample period for Type I neurons (*p* = 3.8 × 10^-3^, *t_4_* = 6.0, paired *t*-test).

### Trial-by-trial changes in neuronal delay activity and performance change

Further, we analyzed the trial-by-trial changes in the delay activity of each subtype, i.e., Type I–III, of the increased type neurons during neurofeedback because the decreased type neurons did not have enough statistical power due to the small population size (<10 for the neurons with neurofeedback). For this analysis, we used the neurons for which ≥15 trials of each stimulus were tested during the neurofeedback blocks (Type I: *n* = 51; Type II: *n* = 15; Type III: *n* = 12). All neuronal types increased their delay activity from the first trial for both sample stimuli (**Fig. 5a–c**). These delay-length modifications of Type I and II neurons mostly continued throughout the trials (**Fig. 5a,b**). Notably, for Type III neurons, the differences between the stimuli in delay activity increased as the trial proceeded, which was formed by a decrease in delay activity for the non-preferred stimulus at the population level (stimulus factor: *p =* 9.3 × 10^-3^, *F*_[1, 280]_ = 1.79; interaction: *p =* 6.6 × 10^-2^, *F*_[14, 280]_ = 1.65, repeated two-way ANOVA with the trial and stimulus factors; **Fig. 5c**). To determine when the differentiation between the preferred and non-preferred stimuli started, we compared delay activity between the stimuli in each trial during the neurofeedback blocks. The differential activity began on the eighth trial after neurofeedback started (*p* < 0.05, *t*-test in each trial; **Fig. 5c**). Among the 12 Type III neurons tested, two-thirds (8/12) did not show differential delay activity between the stimuli in the constant blocks (*n.s.*, paired *t*-test). These results indicate that Type III neurons developed a preference for a specific stimulus throughout neurofeedback.

**Figure 5.**
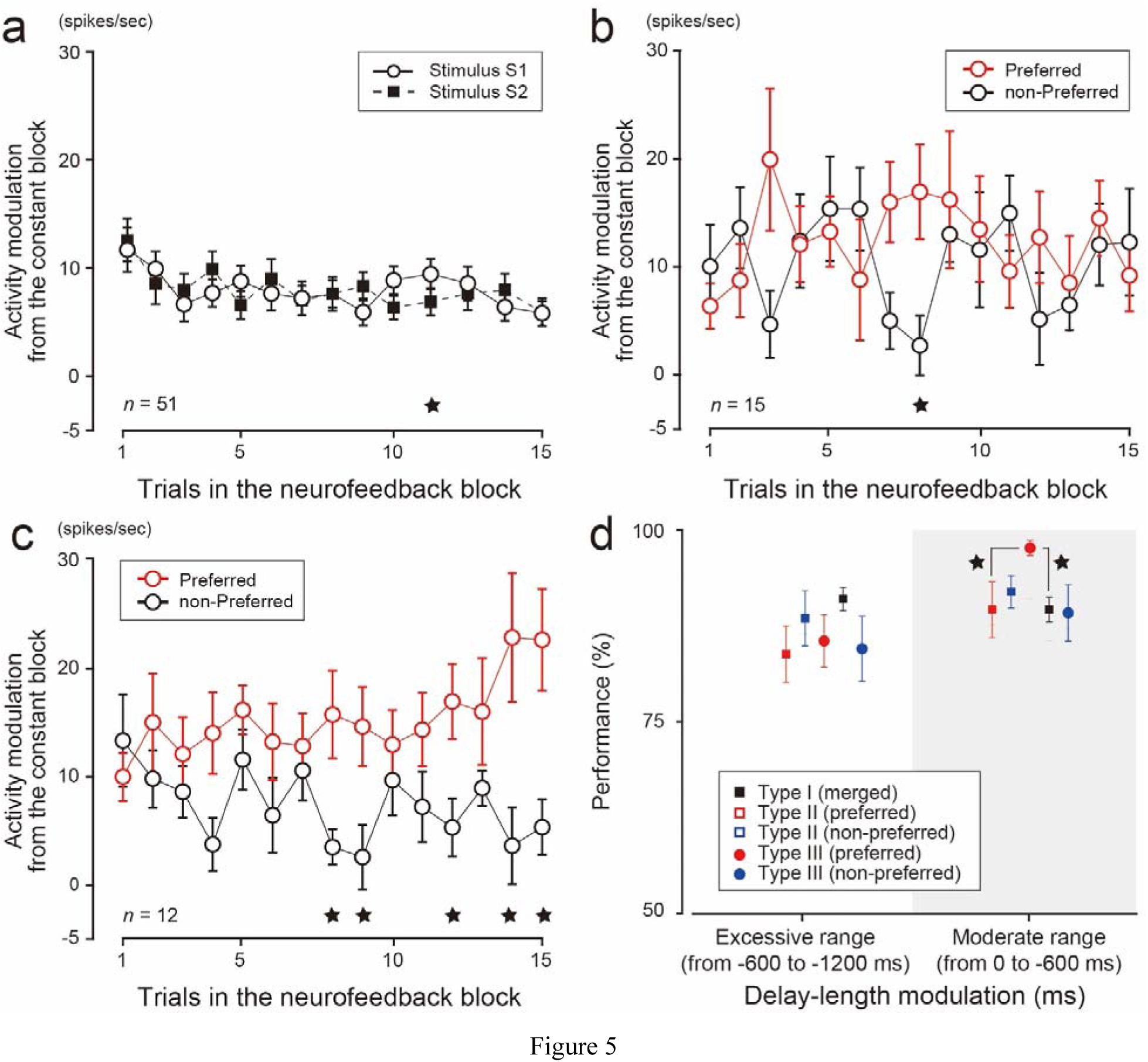
Trial development of neuronal activity for each type (**a–c**) and correct performance rates of each neuronal type in the excessive and moderate ranges of delay-length modulation (**d**). **a–c**. Activity modulation of Type I (**a**), Type II (**b**), and Type III (**c**) neurons during the delay period in the first 15 trials in the neurofeedback blocks. For Type I neurons, black squares and white circles illustrate activity modulation for the stimuli in the neurofeedback blocks compared with the constant blocks (**a**). For the other two types of neurons, lines display activity modulation for the preferred (red circles) and non-preferred (black circles) stimuli (**b** and **c**). Black stars denote significant differences in firing rate between the stimuli during the delay period in the trial (*p* < 0.05, *t*-test). For Type III neurons, a divergence of neuronal modulation during the delay period between the preferred vs. non-preferred stimuli developed throughout the trials (**c**). **d.** Correct performance was retained with moderate delay-length modulation (- 600–0 ms) but not out of this range, in the excessive epoch (from -1200 to -600 ms), when the preferred stimulus of Type III neurons was tested under neurofeedback. Note that sample sizes depend on the delay-length modulation epochs (excessive vs. moderate) and that the positions of neuronal types do not indicate exact delay-length modulation. Stars indicate statistical significance (*p* < 0.05, *t*-test). Shaded area represents the epoch of moderate delay-length modulation. Error bars indicate the standard error of the mean.

We next assessed how the delay activity of each neuronal type was correlated with the correct performance rate depending on delay-length modulation during the neurofeedback blocks. For this analysis, the correct choice rate was computed separately for Type II and III neurons in two analysis epochs (<-600 ms and -600–0 ms), but the data for both stimuli were merged for Type I neurons because there was no significant difference in all epochs. The correct choice rate of the preferred stimulus for Type III neurons was significantly higher than that of the non-preferred stimulus (*p* = 3.5 × 10^-2^, *t*_9_ = 2.5, paired *t*-test) and that of Type I neurons (*p* = 9.2 × 10^-6^, *t*_55_ = 4.9, Welch’s *t*-test) when delay-length modulation was within - 600–0 ms (**Fig. 5d**). In contrast, the correct choice rate was similarly reduced when delay-length modulation was not in this range (<-600 ms). These results suggest that Type III neurons contributed to the high performance rate for a specific stimulus under neurofeedback in the moderate range of delay-length modulation (-600–0 ms).

### Modulation of simultaneously recorded irrelevant neurons

We also classified the irrelevant neurons into Type I–III neurons. Among the increased type irrelevant neurons, we obtained 20 Type I neurons (20/30, 66.7%), 6 Type II neurons (6/30, 20%), and 4 Type III neurons (4/30, 13.3%) (**Fig. 6**). This finding indicates that similar proportions of Type I–III neurons without neurofeedback linkage to those with neurofeedback linkage were recruited in the net activity of the LPFC. We further analyzed what types of irrelevant neurons were observed when a specific type of neuron with neurofeedback was modulated. The analysis revealed that specific combinations of neuronal types between the neurofeedback-given neurons and the irrelevant neurons were not recruited by neurofeedback (**Table S1**). These results strengthened our hypothesis that neurofeedback reconfigured net activity in the LPFC.

**Figure 6.**
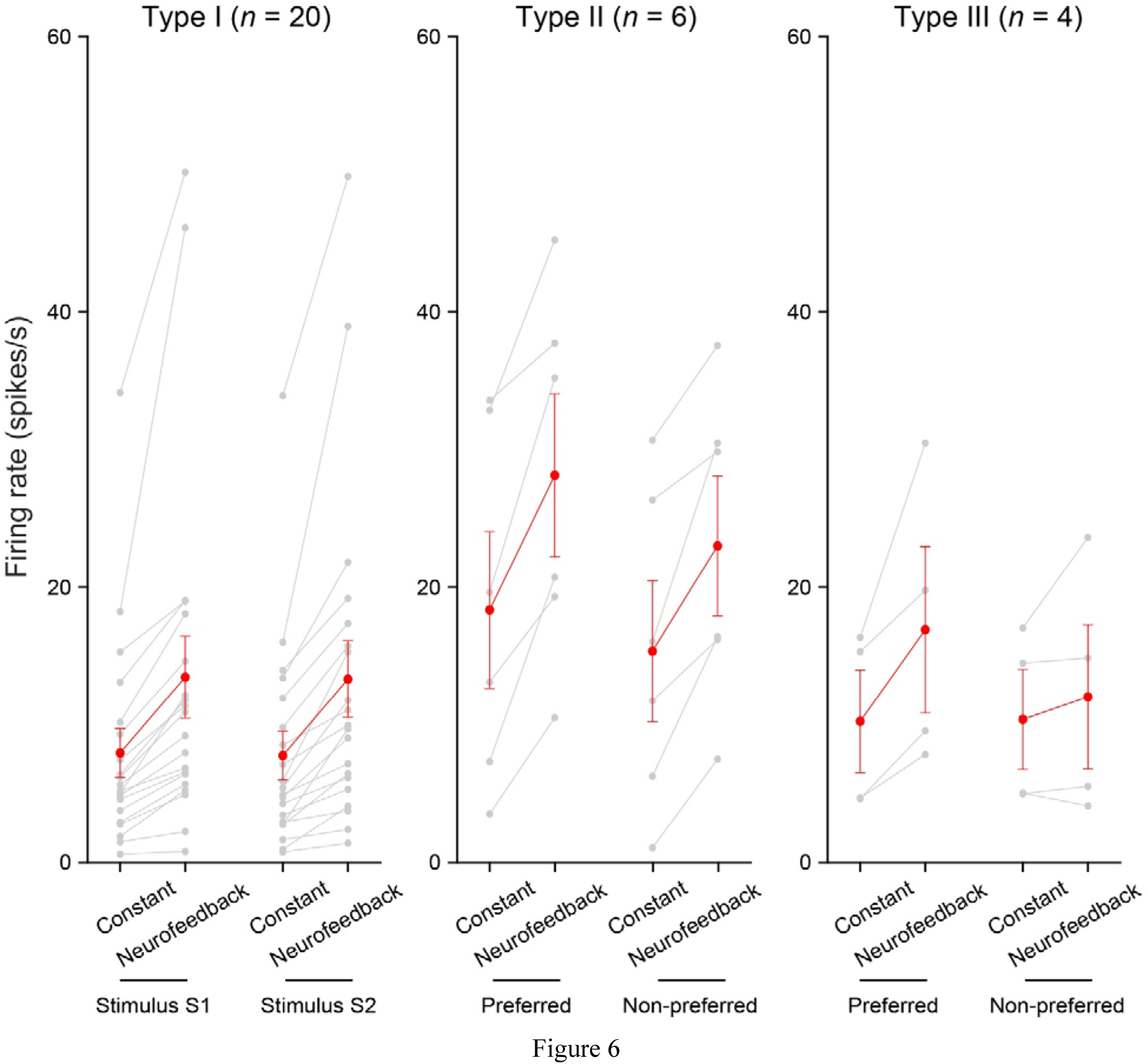
Firing rate and number of irrelevant neuronal types. Type I (*n* = 20, *left*), Type II (*n* = 6, *middle*), and Type III (*n* = 4, *right*) neurons. Irrelevant neurons were recorded simultaneously without neurofeedback of their neuronal activity. Error bars depict the standard error of the mean.

## Discussion

Despite there being no necessity to shorten the delay length in the neurofeedback blocks, both monkeys did so by effectively increasing the delay activity of most neurofeedback-given neurons (**Figs. 3 and 4**). This indicates that the monkeys were strongly motivated to shorten the delay length, as a large body of work suggests that organisms prefer earlier consequences and deterministic situations rather than delayed, uncertain, or ambiguous situations (Hayden, 2016; Kim et al., 2009; Sakurai, 2001). However, such motivation did not simply contribute to correct performance in our neurofeedback WM task. Excessive increased neuronal modulation induced more choice errors and decreased WM performance (**Fig. 2a,c**). In contrast, decreased neuronal modulation functioned oppositely to lengthen the delay period against the animals’ motivation and intention (**Fig. S1**). The decrease in correct performance was recovered in the yoked blocks despite having the same delay-length variance as in the neurofeedback blocks. This finding also suggests that neurofeedback, not delay-length variance, was involved in the decreased performance and reconfiguration of neuronal activity in the LPFC.

Few neurofeedback studies using the unit activity of LPFC neurons have been reported (Kobayashi et al., 2010); however, the consequences of neurofeedback were directly linked with rewards in these studies. In the present study, we devised a new task in which the delay period could be shortened by increasing the delay activity of single LPFC neurons, which was not linked directly to the outcome. Therefore, our neurofeedback was likely associated with WM rather than reward-related functions. Using this neurofeedback task, we successfully conditioned the delay activity of LPFC neurons and tracked the changes of neuronal activity among the blocks. We identified generally increased neuronal types (Type I, 54/82; Type II, 15/82) and a specific stimulus-preferring neuronal type (Type III, 13/82) in which stimulus preference emerged during neurofeedback without a decrease in performance. Our identification of irrelevant neurons suggests that neurofeedback recruited all types of neurons in the LPFC and reconfigured net activity, such as the proportion of Type I–III neurons and the decreased type neurons or their activity balance, to adapt to situational changes. Thus, we demonstrated that neurofeedback using LPFC unit activity effectively modified the activity of LPFC neurons to adapt to a new situation.

Under neurofeedback, Type I–III neurons conceivably played different roles. Type I neurons may largely reflect arousal level. Indeed, these neurons increased their activity not only during the delay period but also in the fixation and sample periods in the neurofeedback blocks (**Fig. 4b**). Although the enhanced activity predominantly during the delay period might facilitate the processing function of WM, its excessive modulation likely lowered the signal-to-noise ratio of relevant information to irrelevant information and would decrease the performance rate. Some electrophysiological studies using non-human primates support this notion (Qi et al., 2011, 2012) by demonstrating that more PFC neurons are recruited during stimulus presentation and delay period in WM tasks, but the mean selectivity of individual neurons to discriminate stimuli decreases. These findings suggest that balancing the recruited neuron population and representation activity of relevant information would be crucial to optimize the signal-to-noise ratio for WM. Similarly, neurofeedback of Type II neurons might induce a lower signal-to-noise ratio in net activity for WM under neurofeedback, although they have the potential that robustly contribute to the correct choice by maintaining differential delay activity through situational changes. Although it remains unclear why only Type III neurons could resist decrease in correct performance by neurofeedback, the emergence of preference may reflect adaptation to the situational changes during the neurofeedback blocks.

This view is consistent with the framework of adaptive coding to environments, which contains concepts of WM and selective attention and control (Duncan, 2001; Gdalyahu et al., 2012; Watanabe and Sakagami, 2007). It hypothesizes that PFC neurons simultaneously encode different types of information, i.e., multiplex, related to current concerns and behavior, such as stimuli, responses, memory, and reward, by inactivating and activating neurons in a situation-dependent manner. The decreased-type neurons, albeit representing a small population in the LPFC, might play a critical role in regulating the activity of LPFC neurons, even against the task demand and the animal’s will. Our identification of these neuronal types may be proof of adaptive coding in the LPFC.

The emergence of preference in WM function, as observed for the activity of Type III neurons (**Fig. 4d**), is often described by recurrent neural network models using excitatory and inhibitory units. Such models can successfully trace the delay activity of PFC neurons and generate a preference or tuning to a specific stimulus through training in WM tasks (Liu et al., 2021; Wang, 2002, 2008). Indeed, cortical circuits, including the LPFC, have recurrent connections (Camperi and Wang, 1998; Goldman-Rakic, 1995). Despite having a different time scale from the models in these studies, our findings of the emergence of preference in Type III neurons and increased and decreased type neurons, i.e., Type I and II neurons, were consistent with their results. The use of recurrent neural network models may also subserve to study the relationships between behaviors and unit activity, mainly to understand the neuronal characteristics of Type III neurons, decreased type neurons, and the excitation-inhibition balance in future studies.

On the basis of the neuronal modulation induced by neurofeedback, net reconfiguration conceivably occurred not only in the LPFC but also in other brain regions such as the ventral striatum (Pereira et al., 2023), parietal cortices (Garrison et al., 2021), and medio-fronto cortices including the dorsolateral PFC, premotor area, and supplemental motor area (Newton et al., 2011) during WM tasks. In fact, the reorganization of these networks has been observed under neurofeedback training in a WM task, although this training was provided solely to dorsolateral PFC activity (Zhang et al., 2015). That study also reported increases and decreases of activity in the targeted brain region through training and neurofeedback, similarly to our case. These studies and our findings also suggest that the balance of excitation and inhibition in these brain networks including the LPFC may play a critical role in maintaining or increasing WM performance.

The observed decrease in correct performance is consistent with the results of previous studies showing that performing dual tasks interferes with the performance of each task (Watanabe and Funahashi, 2018). In such dual tasks, the ability of prefrontal neurons to represent task-relevant information decreases as the demand for the concurrent counterpart task increases (Watanabe and Funahashi, 2014). These studies suggest that the decrease in performance in our task could be due to the overload of multitasking cognitive control by neurofeedback (Cole et al., 2013; Cole et al., 2012). In multitasking cognitive control situations, the same LPFC neural population is simultaneously recruited and overloaded by multiple tasks. Consequently, the limited neural resources in the LPFC are shared, which impairs the performance of the targeted task. In our case, during the delay period in the neurofeedback blocks, the monkeys might face dual cognitive operations, i.e., retaining the sample stimulus in WM and shortening the delay period by increasing the delay activity. The increased activity of Type I–III neurons might reduce the neural resources available for WM and decrease performance. In contrast, in the yoked blocks, in which the association between delay activity and length was dissected, the resource of delay activity was released, and performance recovered. Thus, our findings might demonstrate that LPFC neurons adapted to the new neurofeedback situation under multitasking with the constraint of finite resources. Although we could not collect enough unit data in the yoked blocks in the present study, chronic recording spike data would allow us to address this question. Another possibility is insufficient adaptation time for the animals. Previous work that successfully improved the performance of WM tasks included a training period using neurofeedback for adaptation to the neurofeedback condition, whereas we provided neurofeedback immediately after the constant block. Such a manner of applying neurofeedback might not supply the animals with sufficient time for a better or optimal reorganization of the local network in the LPFC or the WM-related network, resulting in the increase of choice errors and similar saccadic latency at choice errors with short delay-length modulation in the neurofeedback blocks compared to those in the constant blocks. In this study, we used neurofeedback of single-unit activity and revealed the net-activity modulation. It is possible that neurofeedback of net LPFC activity can manipulate and improve WM performance more precisely, as observed in imaging studies using neurofeedback approaches with various computational algorithms, e.g., decoded neurofeedback and real-time functional magnetic resonance imaging techniques, in which brain activity is led to a target state to improve the learning of subthreshold perception (Shibata et al., 2011) and motor-related dysexecutive behavior clinically (Mazrooyisebdani et al., 2018; Mohanty et al., 2018; Papoutsi et al., 2018). A further step is to directly and causally manipulate the net state of LPFC neuronal activity in WM tasks and identify how these three neuronal types and the decreased type contribute to WM performance.

In conclusion, we identified different neuronal types, such as the increased and decreased types of Type I–III neurons during WM under neurofeedback of LPFC unit activity. We showed the effectiveness of this technique to track and monitor real-time neuronal modulation parallel to situational changes. Further, we demonstrated the significant potential of neurofeedback using unit activity to investigate information processing in the brain.

## Methods

### General

All experimental surgical protocols were approved by the Animal Care and Use Committee at Tamagawa University and were in accordance with the US National Institutes of Health Guide for the Care and Use of Laboratory Animals. Two male Japanese monkeys (*Macaca fuscata*, monkey M: 6.5 kg and monkey S: 7.0 kg) were the subjects in the present study. Each animal was mildly fluid deprived during the task-training and experimental periods. In the experiment, the monkey sat in a primate chair inside a sound-attenuated room with its head fixed. Visual stimuli were presented on a 21-inch computer monitor (FlexScan T966, EIZO, Ishikawa, Japan) placed 90.0 cm in front of the monkey at eye level. Single-unit activity around the principal sulcus area in the left hemispheres was recorded by using a tungsten electrode (diameter, 0.25 mm; 0.5–2 MΩ; Frederick Haer, Bowdoinham, ME, USA). The recording sites were determined by reference to magnetic resonance images (**Fig. 1a**). Eye position was monitored at 240 Hz by an infrared eye-tracking system (resolution, 0.25° in visual angle; EYE-TRAC6; Applied Science Laboratories, Bedford, MA, USA). The Tempo system (Reflective Computing, Olympia, WA, USA) was used to control the experiment and data acquisition with 1-ms temporal resolution.

### Surgical procedures

Under sterile conditions, the monkeys were surgically prepared under ketamine HCl (10 mg/kg, i.m.) and pentobarbital-induced anesthesia (Nembutal, 25 mg/kg, i.p.; note that this study was conducted during the lawful allowance of pentobarbital-induced anesthesia.) After exposing the skull, acrylic screws were installed to fasten dental acrylic head implants to the skull. A hollow rod (diameter, 15 mm) was attached to the skull with dental acrylic for head fixation. A recording chamber (30 × 40 mm) was positioned stereotaxically over the LPFC (left hemisphere in both animals) to access the ventral and dorsal areas around the principal sulcus (**Fig. 1a**) and secured with dental acrylic. Antibiotics (ampicillin, i.m., total 1 g/day) and analgesics (diclofenac sodium, 6.25 mg) were administered on the day of surgery and for seven and three days postoperatively, respectively. Craniotomy was performed after the monkeys had learned the constant tasks well (performance >85%).

### DMPST

We trained the animals to perform a visual DMPST. A trial started with the presentation of a fixation point at the center of the monitor screen (**Fig. 1b**). The animal was required to fixate on it for at least 700 ms (fixation period) within a diameter of 3°. A sample stimulus was then presented for 500 ms at the center of the screen (sample period), followed by a delay (delay period). At the beginning of the delay period, the fixation color changed to red, and two small rectangles appeared at 3° below the stimulus. Each rectangle moved horizontally, away from the other, toward the right and left edges of the screen at the same speed. When both rectangles reached the edges, they disappeared, indicating the end of the delay period. Two test stimuli then appeared at 5° right and left from the screen center. The animal was required to choose one of them by making a saccadic eye movement in 500 ms and hold it for at least 300 ms within a 3° diameter. A water reward (0.1 mL per trial) was delivered when the animal chose the paired associate correctly (correct trials). In the experiment, two sample stimuli (S1 and S2) and two test stimuli (T1 and T2) were used; S1 was paired with T1, and S2 was paired with T2 (**Fig. 1c**). One of the sample stimuli was used pseudorandomly in a trial. The right-left positions of the test stimuli were also determined pseudorandomly. No reward was delivered when the animal made eye-fixation breaks, eye position outside of a 3° visual window (fixation-break error trials), or wrong choices by choosing the unpaired target (choice-error trials). Since eye fixation was required during the fixation, sample, and delay periods, breaking fixation during these periods resulted in trial termination with a time-out period of 5 s in addition to the inter-trial interval (3–4 s). After a fixation-break error, the same stimulus and test stimuli were presented in the subsequent trial (correction trials). In the correction trials, half the reward (0.05 mL) was delivered when the animal correctly chose the paired associate. The correction trials were excluded from the analysis.

### Constant, neurofeedback, and yoked blocks

In the neurofeedback blocks, rectangle movement was associated with the delay activity of the recorded neurons, i.e., the spike firing rate during the delay period drove the movement of the rectangles. The step size of rectangle movement per spike was determined as a value of the mean firing rate during the delay period in the preceding constant blocks divided by the half-screen size in pixels. For example, when the mean firing rate during the delay period in the constant blocks was 20 spikes/s, and the half-screen size was 540 pixels, each step of rectangle movement was 27 pixels/spike. When the value was indivisible, it was rounded down to the nearest decimal. When the firing pulses of the recorded neurons reached the spike number of the mean firing rate measured in the constant blocks, the rectangles reached the screen edges. This linkage of neuronal activity with rectangle movement allowed the animals to shorten the delay length by increasing the firing rate during the delay period and obtain an earlier opportunity of target choice. The more the recording neuron fired during the delay period, the quicker the rectangles moved toward the screen edges (**Fig. 1b,d**). Conversely, the delay period would be lengthened when the recorded neuron fired less than in the constant blocks (**Fig. 1b,d**). Consequently, delay length could vary across trials in the neurofeedback blocks. A pair of the constant and neurofeedback blocks was referred to as a session. A single session, i.e., one constant block and one neurofeedback block, was tested for each neuron. In 95 of 125 sessions, a yoked block was additionally provided for the animals (15 sessions for monkey M and 80 sessions for monkey S). In the yoked blocks, the same sequence of delay lengths and stimuli experienced in the preceding neurofeedback blocks was replayed, but with constant rectangle movement speed. Thus, the rectangles moved at a constant speed during the delay period in the constant and yoked conditions. but not in the neurofeedback blocks. A trial was terminated when delay length was >5 s due to the lower activity of the targeted neurons in the neurofeedback blocks than in the constant blocks. Delay length in the constant blocks was constant in a block and varied among the sessions (>1,000 ms; median length, 1,400 ms; interquartile range, 200 ms) to weaken the temporal prediction of delay length. The color of the fixation point instructed the animal as to which block it was to perform: blue for the constant blocks, yellow for the neurofeedback blocks, and green for the yoked blocks.

### Measurement of eye position and movement

To examine the relationships between neuronal activity and eye movement, we measured the horizontal and vertical eye positions and smoothed them using a Gaussian kernel (σ = 10 ms, width = 3 σ). We calculated the onset, offset, peak velocity, amplitude, and duration of saccadic eye movement using the vector data of the square root (dx^2^ + dy^2^) of eye position at a time resolution of 1 ms. Saccade onset was defined as the first time when the angular velocity of a saccade exceeded 30°/s. Saccade offset was defined as the first time when the angular velocity of a saccade was less than 30°/s after saccade onset. The period and maximum velocity between saccade onset and offset were used as saccade duration and peak velocity, respectively. Saccadic amplitude was defined as the distance in degrees between the saccade onset and offset points.

### Task-related neurons

A neuron was classified as a task-related neuron when it showed a significant difference in its firing rate by repeated ANOVA (*p* < 0.05) during at least one of the five epochs in the constant blocks: pre-sample stimulus epoch (0.5 s before sample stimulus), post-sample stimulus epoch (0.5 s after sample stimulus), delay epoch (0.5–1.0 s after sample stimulus), pre-test stimulus epoch (0.5 s before test stimulus), and post-test stimulus epoch (0.5 s after test stimulus). In all 125 sessions, we used the neurons that exhibited significant modulation in the constant blocks of the task (task-related neurons) for our neurofeedback study.

## Supplementary

**Figure S1.**
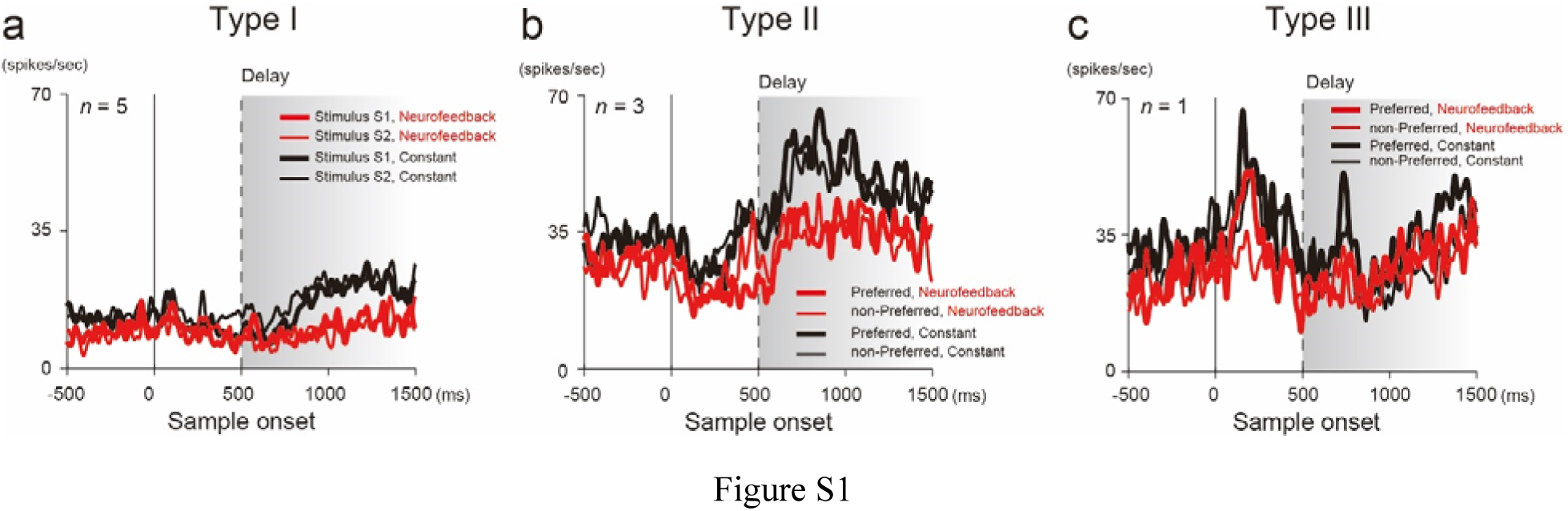
Decreased type activity. **a–c.** Activity of decreased Type I (**a**), II (**b**), and III (**c**) neurons. Vertical dashed lines indicate the start of the delay period. Curves illustrate the mean firing rates of population activity (Type I: *n* = 5; Type II: *n* = 3; Type III: *n* = 1) of the sample stimuli (Type I: stimulus S1, thin curves; stimulus S2, thick curves; Type II and III: non-preferred stimulus, thin curves; preferred stimulus, thick curves). Shaded areas with gradation represent variable delay length. Note that the preferred and non-preferred stimuli were determined dependent on the delay activity in the neurofeedback blocks. Colors represent the constant (black) and neurofeedback (red) blocks. The same convention was used as in Figure 4.

**Table S1.**
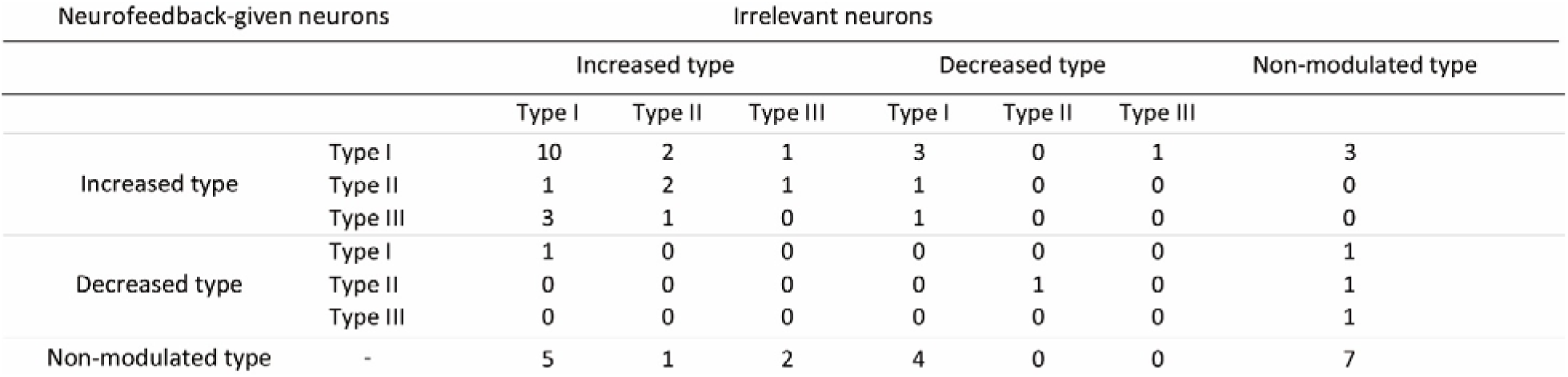
A cross-reference table between the neurofeedback-given neurons and the irrelevant neurons. We counted the number of neurons when the neurofeedback-given neurons and the irrelevant neurons were recorded simultaneously. Note that the irrelevant neurons were not always obtained during the recording of neurofeedback-given neurons. Accordingly, the summation numbers of each neurofeedback-given neuronal type in this table were not equal to the reported numbers for the corresponding neuronal types in the main text.

